# Harnessing genetic diversity in the USDA pea (*Pisum sativum* L.) germplasm collection through genomic prediction

**DOI:** 10.1101/2021.05.07.443173

**Authors:** Md. Abdullah Al Bari, Ping Zheng, Indalecio Viera, Hannah Worral, Stephen Szwiec, Yu Ma, Dorrie Main, Clarice J. Coyne, Rebecca McGee, Nonoy Bandillo

## Abstract

Phenotypic evaluation and efficient utilization of germplasm collections can be time-intensive, laborious, and expensive. However, with the plummeting costs of next-generation sequencing and the addition of genomic selection to the plant breeder’s toolbox, we now can more efficiently tap the genetic diversity within large germplasm collections. In this study, we applied and evaluated genomic selection’s potential to a set of 482 pea accessions – genotyped with 30,600 single nucleotide polymorphic (SNP) markers and phenotyped for seed yield and yield-related components – for enhancing selection of accessions from the USDA Pea Germplasm Collection. Genomic prediction models and several factors affecting predictive ability were evaluated in a series of cross-validation schemes across complex traits. Different genomic prediction models gave similar results, with predictive ability across traits ranging from 0.23 to 0.60, with no model working best across all traits. Increasing the training population size improved the predictive ability of most traits, including seed yield. Predictive abilities increased and reached a plateau with increasing number of markers presumably due to extensive linkage disequilibrium in the pea genome. Accounting for population structure effects did not significantly boost predictive ability, but we observed a slight improvement in seed yield. By applying the best genomic prediction model (e.g., RR-BLUP), we then examined the distribution of genotyped but nonphenotyped accessions and the reliability of genomic estimated breeding values (GEBV). The distribution of GEBV suggested that none of the nonphenotyped accessions were expected to perform outside the range of the phenotyped accessions. Desirable breeding values with higher reliability can be used to identify and screen favorable germplasm accessions. Expanding the training set and incorporating additional orthogonal information (e.g., transcriptomics, proteomics, metabolomics, physiological traits, etc.) into the genomic prediction framework could enhance prediction accuracy.

## Introduction

Pea (*Pisum sativum* L.) is a vitally important pulse crop that provides protein (15.8-32.1%), vitamins, minerals, and fibers. Pea consumption has cardiovascular benefits as it is rich in potassium, folate, and digestible fibers, which promote gut health and prevent certain cancers (Mudryj et al., 2014; Tayeh et al., 2015). Considering the health benefits of pea, the US Department of Agriculture recommends regular pulses consumption, including peas, to promote human health and wellbeing (http://www.choosemyplate.gov/). In 2019, more than 446,000 hectares of edible dry pea were planted with production totaling 1,013,600 tonnes in the USA, making it the fourth-largest legume crop (http://www.fao.org) (USDA, 2020). Growing peas also help maintain soil health and productivity by fixing atmospheric nitrogen (Burstin et al., 2015). Recently, the pea protein has emerged as a frontrunner and showed the most promise in the growing alternative protein market. The Beyond Meat burger is a perfect example of a pea protein product gaining traction in the growing market. About 20-gram protein (17.5%) in each burger comes from pea (https://www.nasdaq.com/articles/heres-why-nows-thetime-to-buy-beyond-meat-stock-2019-12-05). Another product made from pea, Ripptein, is a non-dairy milk product of pea protein that is gaining tremendous interest as an alternative dairy product (https://www.ripplefoods.com/ripptein/). Additionally, peas are gaining attention in the pet food market as it is grain-free and an excellent source of essential amino acids required by cats and dogs (PetfoodIndustry.com) (Facciolongo et al., 2014). As the demand for pea increases, particularly in the growing alternative protein market, genetic diversity expansion is needed to hasten the current rate of genetic gain in pea (Vandemark et al., 2014).

Germplasm collections serve as an essential source of variation for germplasm enhancement that can sustain long-term genetic gain in breeding programs. The USDA *Pisum* collection, held at the Western Regional Plant Introduction Station at Washington State University, is a good starting point to investigate functional genetic variation useful for applied breeding efforts. To date, this collection consists of 6,192 accessions plus a Pea Genetic Stocks collection of 712 accessions. From this collection, the USDA core collection comprised of 504 accessions was assembled to represent ~18% of all USDA pea accessions at the time of construction (Simon and Hannan 1995; Coyne et al., 2005). Subsequently, single-seed descent derived homozygous accessions were developed from a subset of the core to form the ‘Pea Single Plant’-derived (PSP) collection. The PSP is used to facilitate the collection’s genetic analysis (Cheng et al., 2015). The USDA Pea Single Plant Plus Collection (PSPPC) was assembled and included the PSP and Chinese accessions and field, snap and snow peas from US public pea-breeding programs (Holdsworth et al., 2017).

Genomic selection (GS) takes advantage of high-density genomic data that holds a promise to increase the rate of genetic gain (Meuwissen et al., 2001). As genotyping costs have significantly declined relative to current phenotyping costs, GS has become an attractive option as a selection decision tool to evaluate accessions in extensive germplasm collections. A genomic prediction approach could use only genomic data to predict each accession’s breeding value in the collection (Meuwissen et al., 2001; Habier et al., 2007; VanRaden, 2008). The predicted values would significantly increase the value of accessions in germplasm collections by giving breeders a means to identify those favorable accessions meriting their attention from the thousand available accessions in germplasm collections (Longin et al., 2014; Crossa et al., 2016; Jarquin et al., 2016). Several studies used the genomic prediction approach to harness diversity in germplasm collections, including lentil (Haile et al., 2020), soybean (Jarquin et al., 2016), wheat (Crossa et al., 2016), rice (Spindel et al., 2015), sorghum (Yu et al., 2016), maize (Gorjanc et al., 2016), and potato (Bethke et al., 2019). A pea genomic selection study for drought-prone Italian environment revealed increased selection accuracy of pea lines (Annicchiarico et al., 2019; Annicchiarico et al., 2020). To the best of our knowledge, no such studies have been performed using the USDA Pea Germplasm Collection, but a relevant study has been conducted using a diverse pea germplasm set comprised of more than 370 accessions genotyped with a limited number of markers (Burstin et al., 2015; Tayeh et al., 2015).

To date, methods to sample and utilize an extensive genetic resource like germplasm collections remain a challenge. In this study, a genomic prediction approach targeting complex traits, including seed yield and phenology, was evaluated to exploit diversity contained in the USDA Pea Germplasm Collection. No research has been conducted before on genomic prediction for the genetic exploration of the USDA Pea Germplasm Collection. Different cross-validation schemes were used to answer essential questions surrounding the efficient implementation of genomic prediction and selection, including determining best prediction models, optimum population size and number of markers, and impact of accounting population structure into genomic prediction framework. We then examined the distribution of all nonphenotyped accessions using SNP information in the collection by applying genomic prediction models and estimated reliability criteria of genomic estimated breeding values for the assessed traits.

## Material and Methods

### Plant materials

A total of 482 USDA germplasm accession were used in this study, including the Pea Single Plant Plus Collection (Pea PSP) comprised of 292 pea germplasm accessions (Cheng et al., 2015). The USDA Pea Core Collection contains accessions from different parts of the world and represents the entire collection’s morphological, geographic, and taxonomic diversity. These accessions were initially acquired from 64 different countries and are conserved at the Western Regional Plant Introduction Station, USDA, Agricultural Research Service (ARS), Pullman, WA (Cheng et al., 2015).

### DNA extraction, sequencing, SNP calling

Green leaves were collected from seedlings of each accession grown in the greenhouse with the DNeasy 96 Plant Kit (Qiagen, Valencia, CA, USA). Genomic libraries for the Single Plant Plus Collection were prepped at the University of Minnesota Genomics Center (UMGC) using genotyping-by-sequencing (GBS). Four hundred eighty-two (482) dual-indexed GBS libraries were created using restriction enzyme *Ape*KI (Elshire et al., 2011). A NovaSeq S1 1 × 100 Illumina Sequencing System (Illumina Inc., San Diego, CA, USA) was then used to sequence the GBS libraries. Preprocessing was performed by the UMGC that generated the GBS sequence reads. An initial quality check was performed using FastQC (http://www.bioinformatics.babraham.ac.uk/projects/fastqc/). Sequencing adapter remnants were clipped from all raw reads. Reads with final length <50 bases were discarded. The high-quality reads were aligned to the reference genome of *Pisum sativum* (Pulse Crop Database https://www.pulsedb.org/, Kreplak et al., 2019) using the Burrow Wheelers Alignment tool (Version .7.17) (Li and Durbin, 2009) with default alignment parameters, and the alignment data was processed with SAMtools (version 1.10) (Li et al., 2009). Sequence variants, including single and multiple nucleotide polymorphisms (SNPs and MNPs, respectively), were identified using FreeBayes (Version 1.3.2) (Garrison and Marth, 2012). The combined read depth of 10 was used across samples for identifying an alternative allele as a variant, with the minimum base quality filters of 20. The putative SNPs from freeBayes were filtered across the entire population to maintain the SNPs for biallelic with minor allele frequency (MAF) < 5%. The putative SNP discovery resulted in biallelic sites of 380,527 SNP markers. The QUAL estimate was used for estimating the Phred-scaled probability. Sites with a QUAL value less than 20 and more than 80% missing values were removed from the marker matrix. The rest of the markers were further filtered out so that heterozygosity was less than 20%. The filters were applied using VCFtools (version 0.1.16) (Danecek et al., 2011) and in-house Perl scripts. The SNP data were uploaded in a public repository and is available at this link: https://www.ncbi.nlm.nih.gov/sra/PRJNA730349 (Submission ID: SUB9608236). Missing data were imputed using a *k*-nearest neighbor genotype imputation method (Money et al., 2015) implemented in TASSEL (Bradbury et al., 2007). SNP data were converted to a numeric format where 1 denotes homozygous for a major allele, −1 denotes homozygous for an alternate allele, and 0 refers to heterozygous loci. Finally, 30,646 clean, curated SNP markers were identified and used for downstream analyses.

### Phenotyping

Pea germplasm collections (Pea PSP) were planted following augmented design with standard checks (’Hampton,’ ‘Arargorn,’ ‘Columbian,’ and ‘1022’) at the USDA Central Ferry Farm in 2016, 2017, and 2018 (planting dates were March 14, March 28, and April 03, respectively). The central Ferry farm is located at Central Ferry, WA at 46°39’5.1’’N; 117°45’45.4” W, and elevation of 198 m. The Central Ferry farm has a Chard silt loam soil (coarse-loamy, mixed, superactive, mesic Calcic Haploxerolls) and was irrigated with subsurface drip irrigation at 10 min d^−1^. All seeds were treated with fungicides; mefenoxam (13.3 mL a.i. 45 kg-1), fludioxonil (2.4 mL a.i. 45 kg −1), and thiabendazole (82.9 mL a.i.45 kg −1), insecticide; thiamethoxam (14.3 mL a.i. 45 kg −1), and sodium molybdate (16 g 45 kg −1) prior to planting. Thirty seeds were planted per plot; each plot was 152 cm long, having double rows with 30 cm center spacing. The dimensions of each plot were 152 cm × 60 cm. Standard fertilization and cultural practices were used.

The following traits were recorded and are presented in this manuscript. Days to first flowering (DFF) are the number of days from planting to when 10% of the plot’s plants start flowering. The number of seeds per pod (NoSeedsPod) is the number of seeds in each pod. Plant height (PH cm) is defined as when all plants in a plot obtained full maturity and were measured in centimeters from the collar region at soil level to the plants’ top. Pods per plant (PodsPlant) is the number of recorded pods per plant. Days to maturity (DM) referred to physiological maturity when plots were hand-harvested, mechanically threshed, cleaned with a blower, and weighed. Plot weight (PlotWeight, gm) is the weight of each plot in grams after each harvest. Seed yield (kg ha^−1^) is the plot weight converted to seed yield in kg per hectare.

### Phenotypic data analysis

A mixed linear model was used to extract best linear unbiased predictors (BLUPs) for all traits evaluated using the following model:

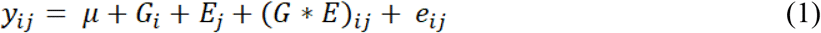

where *y*_*ij*_ is the observed phenotype of *i*^th^ genotypes and *j*^th^ environment which is the number of years, *μ* is the overall mean, *G*_*i*_ is the random genetic effect (*i* is number of genotypes), *E*_*i*_ is the random environments (*j* is number of years),(*G* * *E*)_*ij*_ is the genotype by environment interaction, and *e*_*ij*_ is the residual error.

For the purpose of estimating heritability, we fit the same model above. The heritability in broad sense (*H*^2^) on an entry-mean basis for each assessed trait was calculated to evaluate the quality of trait measurements following the equation (Hallauer et al., 2010):

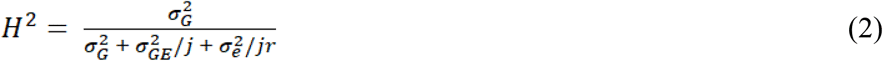

where 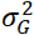 is the genetic variance, 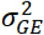 is variance due to the genotype by year interaction, 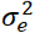 is the error variance, *j* is number of years considered as environments, and *r* is the relative number of occurrences of each genotype in a trial (this is non-replicated trial so harmonic mean of the replicates were used as replicates). We also calculated heritability proposed by (Cullis et al., 2006) implemented in Sommer package in R (Covarrubias-Pazaran, 2016).

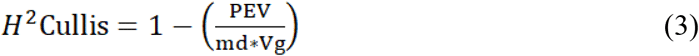

where PEV is the predicted error variance for the genotype, Vg refers to the genotypic variance, md is the mean values from the diagonal of the relationship matrix, which is an identity matrix.

The R package, lme4 (Bates et al., 2015), was used to analyze the data. The trait values derived from the BLUPs were used to measure correlation with the ggcorrplot using ggplot2 package (Wickham 2016). All phenotypic and genomic prediction models were analyzed in the R environment (R Core Team, 2020).

### Genomic selection models

The genomic selection models were fitted as follows:

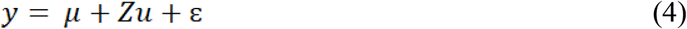

where *y* is a vector of the genotype BLUPs obtained from equation (1), *μ* is the intercept of the model used for the study, *Z* is the SNP marker matrix, *u* is the vector of marker effects, and ɛ is a residual vector.

Five genomic selection methods were used to predict genomic estimated breeding values in respective phenotypes of the assessed traits: ridge regression best linear unbiased prediction approach (RR-BLUP), partial least squares regression model (PLSR), random forest (RF), BayesCpi, and Reproducing Kernel Hilbert Space (RKHS).

The RR-BLUP approach assumes all markers have an equal contribution to the genetic variance. One of the most widely used methods for predicting breeding values is RR-BLUP, comparable to the best linear unbiased predictor (BLUP) used to predict the worth of entries in the context of mixed models (Meuwissen et al., 2001). The RR-BLUP basic frame model is:

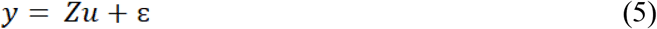

where 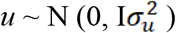 is a vector of marker effects and *Z* is the genotype matrix e.g., {aa,Aa,AA} = {0, 1, 2} for biallelic single nucleotide polymorphisms (SNPs) that relates to phenotype *y* (Endelman, 2011). The RR-BLUP genomic prediction was implemented using the ‘RR-BLUP’ package (Endelman, 2011).

Partial least square regression (PLSR) is a reduction dimension technique that aims to find independent latent components that maximize the covariance between the observed phenotypes and the markers (predictor variables) (Colombani et al., 2012). The number of components (also known as latent variables) should be less than the number of observations to avoid multicollinearity issues and commonly the number of components are chosen by cross validation. PLSR was executed using the ‘pls’ package (Mevik and Wehrens, 2007).

Random forest is a machine learning model for genomic prediction that uses an average of multiple decision trees to determine the predicted values. This regression model was implemented using the ‘randomForest’ package (Breiman, 2001). The number of latent components for PLSR and decision trees for random forest was determined by a five-fold cross-validation to have a minimum prediction error.

BayesCpi was used to verify the influence of distinct genetic architectures of different traits on prediction accuracy. The BayesCpi assumes that each marker has a probability of being included in the model, and this parameter is estimated at each Markov Chain Monte Carlo (MCMC) iteration. The vector of marker effects u is assumed to be a mixture of distributions having the probability π of being null effect and (1-π) of being a realization of a normal distribution, so that, 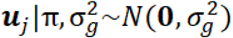. The vector of residual effects was considered as 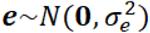. The marker and residual variances were assumed to follow a chi-square distribution 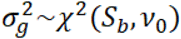 and 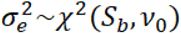, respectively, with *v*_0_ = 5 degrees of freedom as prior and *S*_*b*_ shape parameters assuming a heritability of 0.5 (Pérez and de los Campos 2014).

The last model used was the Reproducing Kernel Hilbert Space (RKHS). The method is a regression where the estimated parameters are a linear function of the basis provided by the reproducing kernel (RK). RKHS considers both additive and non-additive genetic effects (de los Campos et al. 2013). In this work, the multi-kernel approach was used by averaging three kernels with distinct bandwidth values. In this implementation the averaged kernel, 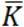 was given by: 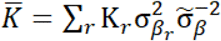, where 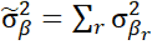. Here r=3 and 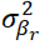 are interpretable as variance parameters associated with each kernel. Therefore, for each r^th^ kernel the proportion of sharing alleles between pairs of individuals (ii′) was given by 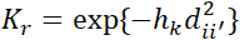, where h_k_ is a bandwidth parameter associated with r^th^ reproducing kernel and 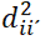 is the genetic distance between individuals i and i′ computed as follows: 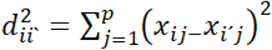, where j=1,…, p markers stated as above. The bandwidth parameter values for the three kernels were h=0.5{1/5,1,5, as suggested by Pérez and de los Campos 2014. Those values were chosen using the rule proposed by de los Campos et al. (2010).

Genomic selection methods RR-BLUP, PLSR, RF were carried out using ‘GSwGBS’ package (Gaynor, 2015) while the BayesianCpi and RKHS were executed with the BGLR package (de los Campos et al., 2010). We calculated each genomic selection model’s predictive ability as the Pearson correlation between the estimated breeding values from model (1) (obtained using the full data set) and those of validation set predicted from the respective model. For that, we used a cross-validation scheme considering 80% of the observations, randomly selected, as training and the remaining 20% as validation set. The process was repeated 20 times for each model. From the predictive ability values, we estimated the confidence interval for this parameter using the bootstrap considering 10000 samples (James et al., 2013).

### Determining optimal training population size

The influence of training population size on predictive ability was evaluated using a validation set comprising 50 randomly selected lines and training sets of variable sizes. The validation set was formed by randomly sampling 50 lines without replacement. The training population of size n was formed sequentially by adding 25 accessions from the remaining accessions such that its size ranged between 50 to 175. We subset the collection into subgroups of 50, 75, 100, 125, 150, and 175 individuals each. The RR-BLUP model was used to predict each trait. This procedure was repeated 20 times, and accuracies of each training population size were averaged across 20 replicates. To predict a particular subpopulation with increasing population size, a similar procedure was followed to using variable training population size ranged from 50 to 175 with an increment of 25.

### Determining optimal marker density

To evaluate the effects of GBS marker selection on predictive ability, we randomly sampled markers five times with the following subset: one thousand (1 K), five thousand (5 K), ten thousand (10 K), fifteen thousand (15 K), twenty thousand (20 K), twenty-five thousand (25 K), and thirty thousand (30 K). A random sampling of SNP was implemented to minimize or avoid any possible biases on sampling towards a particular distribution. Using the RR-BLUP model, a five-fold cross validation approach was used to obtain predictive ability in each marker subset. This procedure was repeated 20 times and predictive ability for each subset of SNP were averaged across 20 replicates.

### Accounting for population structure into the genomic prediction framework

We explored the confounding effect due to population structure on predictive ability. We investigated subpopulation structure on 482 accessions genotyped with 30,600 SNP markers using the ADMIXTURE clustering-based algorithm (Alexander et al., 2009). ADMIXTURE identifies K genetic clusters, where K is specified by the user, from the provided SNP data. For each individual, the ADMIXTURE method estimates the probability of membership to each cluster. An analysis was performed in multiple runs by inputting successive values of K from 2 to 10. The optimal K value was determined using ADMIXTURE’s cross-validation (CV) error values. Based on >60% ancestry, each accession was classified into seven subpopulations (K=7). Accessions within a subpopulation with membership coefficients of <60% were considered admixed. A total of 8 subpopulations were used in this study, including admixed as a separate subpopulation. Principal component (PC) analysis was also conducted to summarize the genetic structure and variation present in the collection.

To account for the effect of population structure, we included the top 10 PCs or, the Q-matrix from ADMIXTURE into the RR-BLUP model and performed five-fold cross-validation repeated 20 times. Alternatively, we also used the subpopulation (SP) designation identified by ADMIXTURE as a factor in the RR-BLUP model. Albeit a smaller population size, we also performed a within-subpopulation prediction. As stated above, a subpopulation was defined based on >60% ancestry cut-off. Only three subpopulations with this cut-off were identified and used: SP5 (N=51), SP7 (N=58), and SP8 (N=41). A leave-one-SP-out was used to predict individuals within the subpopulation with the RR-BLUP model. We also used increasing population sizes to predict specific subpopulation (e.g. SP8) using RR-BLUP model.

### Estimating reliability criteria and predicting unknown phenotypes

Nonphenotyped entries were predicted based on the RR-BLUP model using SNP markers only. The reliability criteria for each of the nonphenotyped lines were then calculated using the formula (Hayes et al., 2009; Clark et al., 2012) as follows:

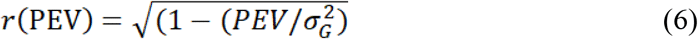

where PEV is the predicted error variance, and 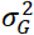 is the genetic variance.

## Results

### Phenotypic heritability and correlation

Recorded DFF had a wide range of variability from 60 to 84 days with a mean of 71 days. The estimated heritability for DFF was 0.90 using equation (2) and 0.80 as per Cullis heritability using equation (3) (**Table 1**). For the number of seeds per pod, the mean was 5.7 with a heritability estimate of 0.84 (*H*^2^_Cullis_=0.66). The heritability for plant height was 0.81 (*H*^2^_Cullis_=0.68), with an average height of 74 cm. Pods per plant had a heritability estimate of 0.50 (*H*^2^_Cullis_=0.27) with a mean of 18 pods per plant and ranged from 15 to 23 pods per plant. DM had a mean of 104 days with an estimated heritability of 0.51 (*H*^2^_Cullis_=0.38). Seed yield per hectare ranged widely from 1734 to 4463 kg ha^−1^ with a mean yield of 2918 kg ha^−1^ and a heritability value of 0.67 (*H*^2^_Cullis_=0.46). The number of pods per plant was highly and positively correlated with seed yield. Correlation estimation also suggested seed yield was positively correlated with plant height (PH), days to maturity (DM), days to first flowering (DFF) (**Supplementary Figure S1**).

**Table 1.** Heritability and summary statistics for seed yield and other agronomic traits

| Trait | Mean | Range | SD | CV(%) | $H^2$ | $H^2_{\text{Cullis}}$ |
| --- | --- | --- | --- | --- | --- | --- |
| DFF (days) | 71 | 60-84 | 4.8 | 6.7 | 0.90 | 0.80 |
| NoSeedsPod (Nos.) | 5.7 | 4.4-6.9 | 0.5 | 8.5 | 0.84 | 0.66 |
| PH (cm) | 74 | 37.6-108.3 | 11.5 | 15.5 | 0.81 | 0.68 |
| PodsPlant (Nos.) | 18 | 15-23 | 1.5 | 8.3 | 0.50 | 0.27 |
| DM (days) | 104 | 99-112 | 2.4 | 2.3 | 0.51 | 0.38 |
| SeedYield (Kg ha<sup>-1</sup>) | 2918 | 1734-4463 | 451 | 15.4 | 0.67 | 0.46 |

DFF is days to first flowering; NoSeedsPod is the number of seeds per pod, PH is plant height, PodsPlant is the number of pods per plant, DM is days to physiological maturity, SeedYield is seed yield per hectare, SD is the standard deviation, CV is coefficient of variance, $H^2$ is heritability in the broad sense.

### Predictive ability of different genomic selection models

No single model consistently performed best across all traits that we evaluated (**Table 2**), however Bayesian model BayesCpi, Reproducing Kernel Hilbert Space (RKHS), and RR-BLUP, in general, tended to generate better results. Roughly the predictive abilities from different models were similar, although slight observed differences were likely due to variations on genetic architecture and the model’s assumptions underlying them. For DFF, the highest predictive ability was obtained from the RR-BLUP (0.60). RR-BLUP, Random Forest (RF), and RKHS models generated the highest predictive ability for pods per plant (0.28). The number of seeds per pod (NoSeedPod) was better predicted by RR-BLUP and Bayes Cpi (0.42). For plant height (PH) highest prediction accuracies were obtained from RF and BayesCpi (0.45). BaysCpi also gave the highest prediction accuracies for DM (0.47). For seed yield, RKHS had slight advantages over other models (0.42). As mentioned above, some differences between the model’s accuracy were only marginal and cannot be a criterion for choosing one model (**Table 2**). For example, among the tested models, the highest difference in predictive accuracy, considering NoSeedsPod, had a magnitude of 0.02, a marginal value. The lack of significant differences among genomic prediction methods can be interpreted as either a good approximation to the optimal model by all methods or there may be a need for further research (Yu et al., 2016). Unless indicated otherwise, the rest of our results focused on findings from the RR-BLUP method.

**Table 2.** Predictive ability of genomic selection models for seed yield and agronomic traits from five genomic selection models

| **Traits** | **RR-BLUP** | **PLSR** | **RF** | **BayesCpi** | **RKHS** |
| --- | --- | --- | --- | --- | --- |
| DFF (days) | 0.60<br>(0.57-0.63) | 0.57<br>(0.53-0.61) | 0.55<br>(0.52-0.58) | 0.59<br>(0.55-0.63) | 0.54<br>(0.5-0.58) |
| NoSeedsPod | 0.42<br>(0.37-0.48) | 0.41<br>(0.36-0.46) | 0.40<br>(0.35-0.45) | 0.42<br>(0.38-0.46) | 0.40<br>(0.34-0.48) |
| PH (cm) | 0.39<br>(0.33-0.44) | 0.42<br>(0.38-0.48) | 0.45<br>(0.4-0.5) | 0.45<br>(0.41-0.48) | 0.43<br>(0.39-0.48) |
| PodsPlant | 0.28<br>(0.22-0.33) | 0.25<br>(0.2-0.31) | 0.28<br>(0.22-0.34) | 0.23<br>(0.17-0.29) | 0.28<br>(0.23-0.34) |
| DM (days) | 0.42<br>(0.36-0.47) | 0.44<br>(0.39-0.5) | 0.41<br>(0.35-0.46) | 0.47<br>(0.43-0.5) | 0.45<br>(0.4-0.48) |
| SeedYield (kg<br>ha-1) | 0.38<br>(0.34-0.42) | 0.31<br>(0.27-0.36) | 0.39<br>(0.35-0.44) | 0.35<br>(0.31-0.39) | 0.42<br>(0.37-0.48) |

DFF is days to first flowering, PH is Plant height in cm, DM is days to physiological maturity; within parentheses are ranges of predictive ability

### Determining the optimal number of individuals

Increasing the training population size led to a slight increase in the predictive ability overall for all traits. Across all traits except days to first flowering and plant height, predictive ability reached a maximum with the largest training population size of N=175 (**Figure 1**). A training population comprised of 50 individuals had the lowest predictive ability across all traits. For days to first flowering, and plant height predictive ability did steadily increase up at N= 150, and prediction ability reached the maximum for most traits at highest training population size with N=175. Regardless of population size, predictive ability was consistently higher for days to first flowering, whereas predictive ability was consistently lower for pods per plant (**Figure 1**). However, while predicting subpopulation 5 highest predictive ability was obtained for plant height (**Supplementary Figure S3**).

**Figure 1.**
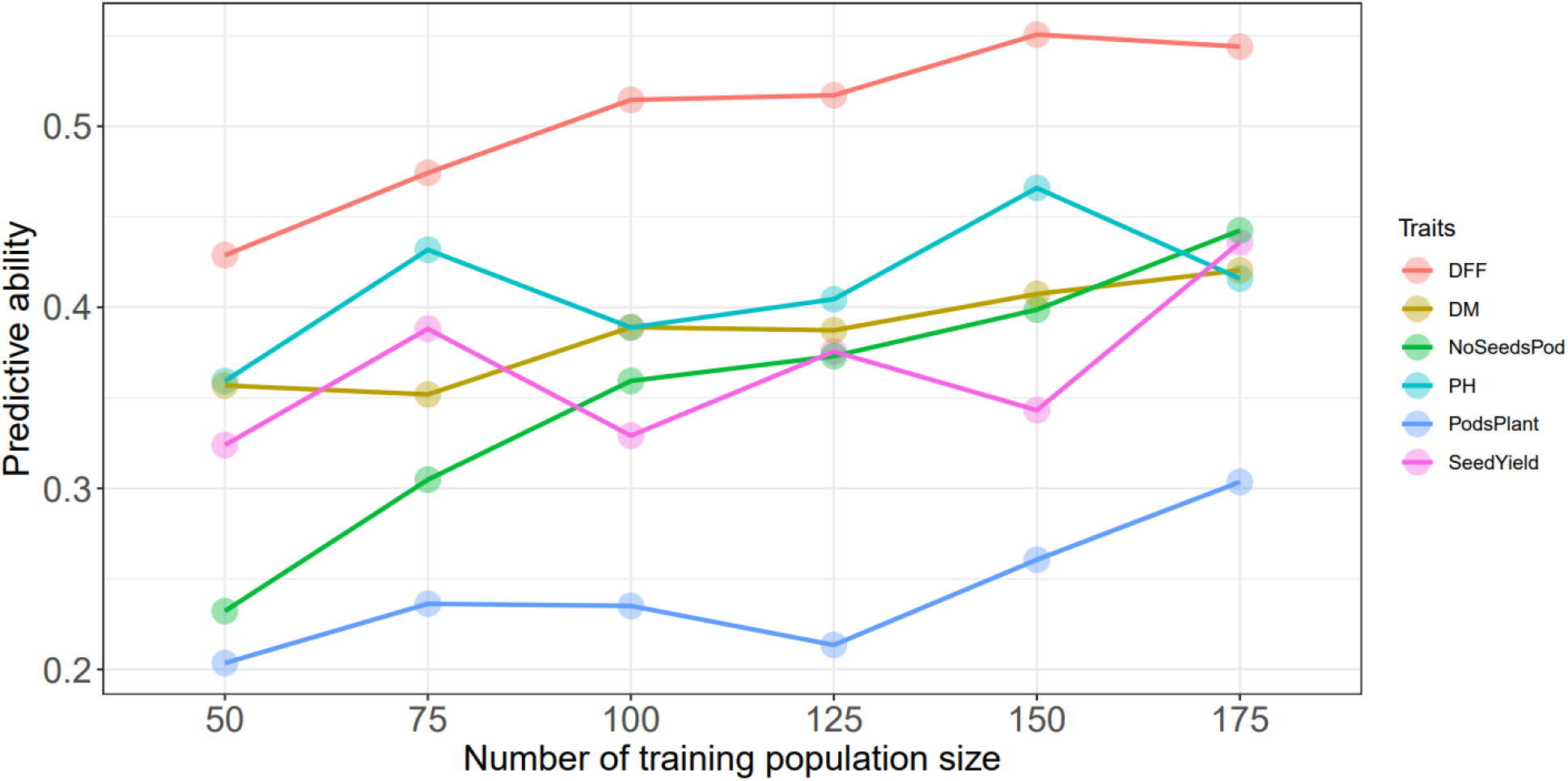
Predictive ability with increasing training population size using RR-BLUP model, DFF is days to first flowering, DM, is days to physiological maturity, NoSeedsPod is number of seeds per pod, PH is plant height in cm, PodsPlant is pods per plant, SeedYield is seed yield in kg ha^−1^

### Determining the optimal marker density

The different marker subsets had insignificant differences on predictive ability for all the traits evaluated in this study. In general, however, predictive abilities were higher between 5K to 15K SNPs and reached a plateau with increasing number of SNP (**Supplementary Figure S2**). For seed yield, plant height, and days to maturity, highest predictive ability were 0.38, 0.39, and 0.42 respectively. The highest predictive ability for DFF was 0.61 using a SNP subset of 15K.

### Accounting for population structure in the genomic prediction model

Population structure explained some portion of the phenotypic variance, ranging from 9-19%, with the highest percentages observed for plant height (19%) and seed yield (17%). Using either ADMIXTURE or PCA to account for the effect due to population structure, we improved the predictive ability. We observed a 6% improvement for days to first flowering and 32% for seed yield compared over models that did not account for population structure.

We also performed within-subpopulation predictions. Presented here are the predictive abilities for subpopulations 5, 7, and 8, as they had at least 40 entries. Subpopulation 8 had the highest predictive ability for days to first flowering (0.68), plant height (0.33), days to maturity (0.43), and seed yield (0.37). The highest predictive abilities for the number of seeds per pod (0.40) and pods per plant (0.12) were obtained from subpopulation 7 (**Table 3**). Notably, predictive ability was generally higher when all germplasm sets or subpopulations were included in the model compared to when predictions were made using a subset of germplasm.

**Table 3.** Predictive ability within and across subpopulations using RR-BLUP and all markers

| Sub pops | DFF | NoSeedsPod | PH | PodsPlant | DM | SeedYield |
| --- | --- | --- | --- | --- | --- | --- |
| Sub pop 5 (51) | 0.27 | 0.26 | 0.08 | -0.01 | 0.02 | 0.18 |
| Sub pop 7 (58) | 0.34 | 0.40 | 0.22 | 0.12 | -0.01 | 0.01 |
| Sub pop 8 (41) | 0.68 | 0.35 | 0.33 | 0.07 | 0.43 | 0.37 |
| SP- | 0.50 | 0.45 | 0.47 | 0.25 | 0.51 | 0.34 |
| SP+ | 0.53 | 0.35 | 0.42 | 0.25 | 0.48 | 0.45 |
| SP PC10 | 0.51 | 0.41 | 0.44 | 0.18 | 0.20 | 0.43 |
| Var exp ($R^2$) | 0.13 | 0.09 | 0.19 | 0.15 | 0.15 | 0.17 |

DFF is days to first flowering, PH is plant height, DM is days to physiological maturity, SP- does not account for population structure, SP+, refers to the population structure addressed in the model, SP PC10 addresses population structure with 10 PC, Var exp ( $R^2$ ) refers the variance explained by population structure after fitting a regression model, within parenthesis represent the number of entries in each subpopulation.

### Predicting genotyped but nonphenotyped accessions

The genomic prediction model was then used to predict nonphenotyped entries based on their SNP information. Based on the distribution of GEBV, none of the predicted phenotypes for nonphenotyped accessions exceeded the top-performing observed phenotypes for seed yield (**Figure 2**). The mean seed yield of predicted entries was 2914 kg/ha, and the mean of observed genotypes 2918 kg/ha were non-significant. The mean of observed and predicted entries were non-significant for the other five traits (Supplementary Table 1). The GEBV for number of pods per plant, number of seeds per pod (**Supplementary Figure S4 and S5**), days to first flowering, and days to maturity all fall within the range of observed phenotypes (Similar Figures not added).

**Figure 2.**
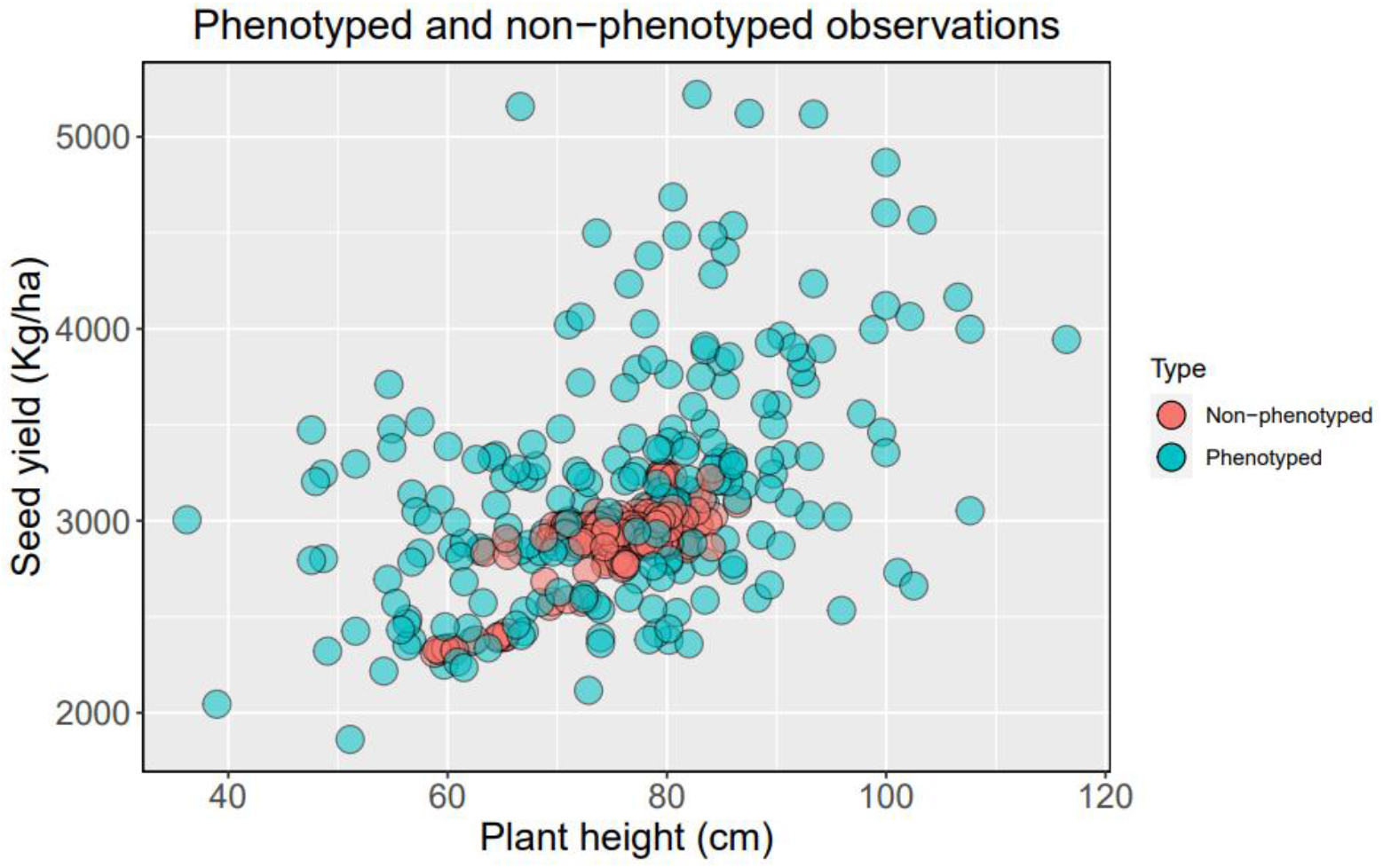
Distribution phenotyped and predicted non-phenotyped accessions of USDA pea germplasm collections for seed yield and plant height

### Reliability estimation

We obtained reliability criteria for all traits, including seed yield and phenology, for 244 nonphenotyped accessions. The average reliability values ranged from 0.30 to 0.35, while the highest values for evaluated traits ranged from 0.75 to 0.78. The higher reliability values were distributed in the top, bottom, and intermediate predicted breeding values (**Supplementary Table S2 to S7**). For seed yield (kg ha^−1^), the highest reliability was obtained from the bottom 50 (**Figure 3**). Higher reliability criteria are primarily distributed among the intermediate and top GEBV for days to first flowering. Predicted intermediate plant height showed the highest reliability, as presented in **Figure 3**.

**Figure 3.**
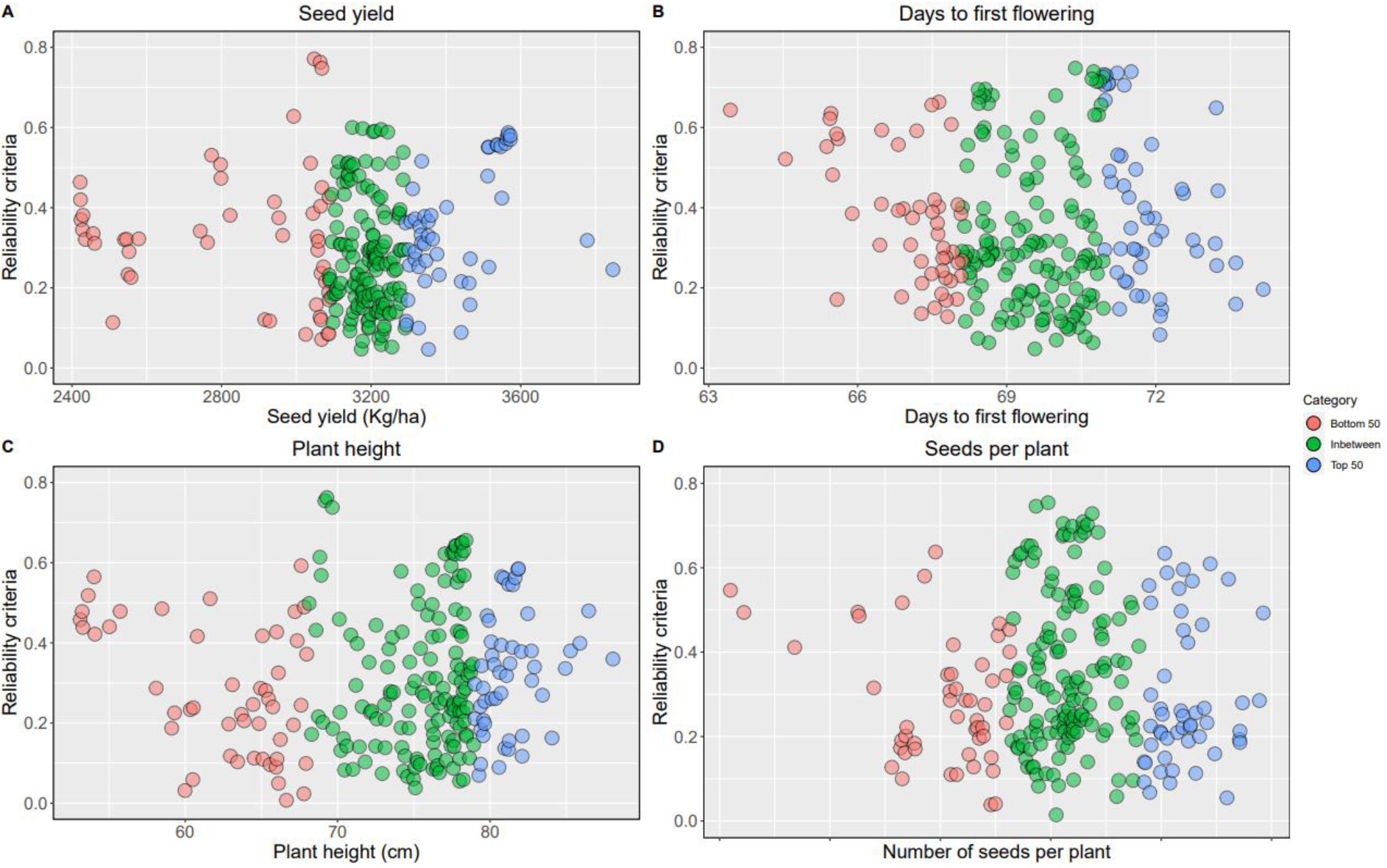
Reliability criteria for nonphenotyped lines: the top 50 of the genomic estimated breeding values are blue, and bottom 50 are in red, intermediates are in green. A. reliability estimates for seed yield (Kg/ha), B. days to first flowering, C. plant height, D. seeds per plant

## Discussion

Widely utilized plant genetic resources collections, such as the USDA pea germplasm collection, hold immense potential as diverse genetic resources to help guard against genetic erosion and serve as unique sources of genetic diversity from which we could enhance genetic gain, boost crop production, and help reduce crop losses due to disease, pests, and abiotic stresses (Crossa et al., 2017; Holdsworth et al., 2017; Jarquin et al., 2016; Mascher et al., 2019). As the costs associated with genotyping on a broader and more accurate scale continue to decrease, opportunities increase to utilize these collections in plant breeding. Relying on phenotypic evaluation alone can be costly, rigorous, and time-intensive. However, by incorporating high-density marker coverage and efficient computational algorithms, we can better realize the potential for utilizing these germplasm stocks by reducing the time and cost associated with their evaluation (Yu et al., 2016; H. Li et al., 2018; Yu et al., 2020). In this study, we evaluated the potential of genotyping-by-sequencing derived SNP for genomic prediction. We found that it holds promises for extracting useful diversity from germplasm collections for applied breeding efforts.

In this study, predictive ability was generally similar among methods, and there was no single model that worked across traits, consistent with results obtained by other authors (Burstin et al., 2015; Spindel et al., 2015; Yu et al., 2016; Azodi et al., 2019). For example, considering only the punctual estimates, RR-BLUP model was the best for DFF, however for PH, DM, and seed yield, the best models were BayesCpi and RF, BayesCpi and RKHS, respectively. In recent work, Azodi et al., (2019) compared 12 models (6 linear and 6 non-linear) considering 3 traits in 6 different plant species, and they did not find any best algorithm for all traits across all species. Newer statistical methods are expected to boost prediction accuracy; however, the biological complexity and unique genetic architecture of traits can be regarded as the root cause for getting zero or slight improvement on prediction accuracy (Yu et al., 2020; Valluru et al., 2019). As data collection accelerates in at different levels of biological organization (Kremling et al., 2019), genomic prediction models will expand and nonparametric models, including machine learning, may play an essential role for boosting prediction accuracy (Azodi et al., 2019; Yu et al., 2020).

A related work in pea has been published but only based on a limited number of markers (Burstin et al., 2015). This work assessed genomic prediction models in a diverse collection of 373 pea accessions with 331SNPs markers and found no single best model across traits, which is consistent with our findings. In this work, the authors reported that traits with higher heritability, such as thousand seed weight and flowering date, had higher prediction accuracy. We also verified DFF as having the highest heritability and predictive accuracies through all the models. Interestingly, yield components like the number of seeds per pod and pods per plant showed lower predictive accuracy, regardless of prediction models used. Consistent with our results, Burstin et al. (2015) also found yield components (seed number per plant) as having lower predictive accuracy and higher standard deviation for prediction. These traits are highly complex and largely influenced by the environment.

The predictive ability increased for all traits except plant height when we increased the model’s training population size, suggesting that adding more entries in the study can boost predictive ability. By accounting population structure into genomic prediction framework, we observed an improved prediction accuracy for some traits – seed yield and DFF – but not others. Although the population structure explained 9-19% of the phenotypic variance, we cannot fully and conclusively answer the effect of population structure in prediction accuracy due to smaller population size. In addition, accounting for the relatedness among individuals in the training and testing sets can potentially boost prediction accuracy (Lorenz and Smith, 2015; Rutkoshi et al., 2015; Riedelsheimer et al., 2013); it was outside the scope of this research but deserves further study. Adding more environments (year-by-location combination) can also potentially improve prediction accuracy using genomic prediction frameworks that account for genotype-by-environment interactions and/or phenotypic plasticity (Jarquin et al, 2014; Crossa et al., 2017; X. Li et al., 2018; Guo et al., 2020). In general, we observed that predictive ability slightly increased and plateaued after reaching certain subset of SNPs. Such a plateau on prediction ability maybe due to overfitting of models (Norman et al., 2018; Hickey et al., 2014), presumably due to extensive linkage disequilibrium in the pea genome (Kreplak et al., 2019).

Previous studies have indicated the importance of considering reliability values when using predictive ability values to select genotypes (Yu et al., 2016). We found higher reliability estimates were spread across all GEBVs rather than clustering around higher or lower extreme of GEBVs. Those accessions with top predicted values and high-reliability estimates maybe selected as candidate parents for increasing seed yield and/or germplasm enhancement. However, for a trait such as days to flowering in pea, even low or intermediate predicted values maybe suitable candidates when paired with high-reliability values. We found the means of GEBV for nonphenotyped entries were non-significantly different with phenotyped accessions, and almost none of nonphenotyped accessions were expected to exceed seed yield of phenotyped accessions. Several accessions in the USDA pea germplasm collection can be readily incorporated into breeding programs for germplasm enhancement by incorporating above-average accessions with high or moderately high-reliability values (Yu et al., 2020).

## Conclusions and Research Directions

The research findings demonstrated that the wealth of genetic diversity available in a germplasm collection could be assessed efficiently and quickly using genomic prediction to identify valuable germplasm accessions that can be used for applied breeding efforts. With the integration of more orthogonal information (e.g., expression, metabolomics, proteomics, etc.) into genomic prediction framework (Kremling et al., 2019; Valluru et al., 2019) coupled with the implementation of more complex genomic selection models like a multivariate genomic selection approach (Rutkoski et al., 2015), we can considerably enhance predictive ability. This research framework could greatly contribute to help discover and extract useful diversity targeting high-value quality traits such as protein and mineral concentrations from a large germplasm collection in the future.

## Author Contributions

NBB, CJC, and MAB conceived and designed the manuscript. CJC, DM, and RMcG designed and executed the field and genotyping experiments. YM and PZ performed DNA extraction, constructed the library, and called SNPs. MAB, IV, and SS analyzed data, curated SNPs, and ran genomic selection models. NBB oversaw statistical analyses. MAB, HW, IV, and NBB wrote and edited the overall manuscript. All authors edited, reviewed, and approved the manuscript.

## Acknowledgments

The authors would like to acknowledge the funding provided by USDA Plant Genetic Resource Evaluation Grant for the GS analysis, USA Dry Pea and Lentil Council Research Committee for the field phenotyping, and USDA ARS Pulse Crop Health Initiative for the SNP genotyping and support from USDA ARS Project: 5348-21000-017-00D (CJC), and 5348-21000-024-00D (RJM). We also acknowledge the support from USDA-NIFA (Hatch Project ND01513 for NBB) and the North Dakota Department of Agriculture through the Specialty Crop Block Grant Program (19-429). Technical assistance from Britton Bourland, Lydia Fields, Kurt Tetrick, and Jennifer Morris gratefully acknowledged. This work used resources of the Center for Computationally Assisted Science and Technology (CCAST) at North Dakota State University, Fargo, ND, USA which were made possible in part by NSF MRI Award No. 2019077.

---

**Supplementary Figure S1.**
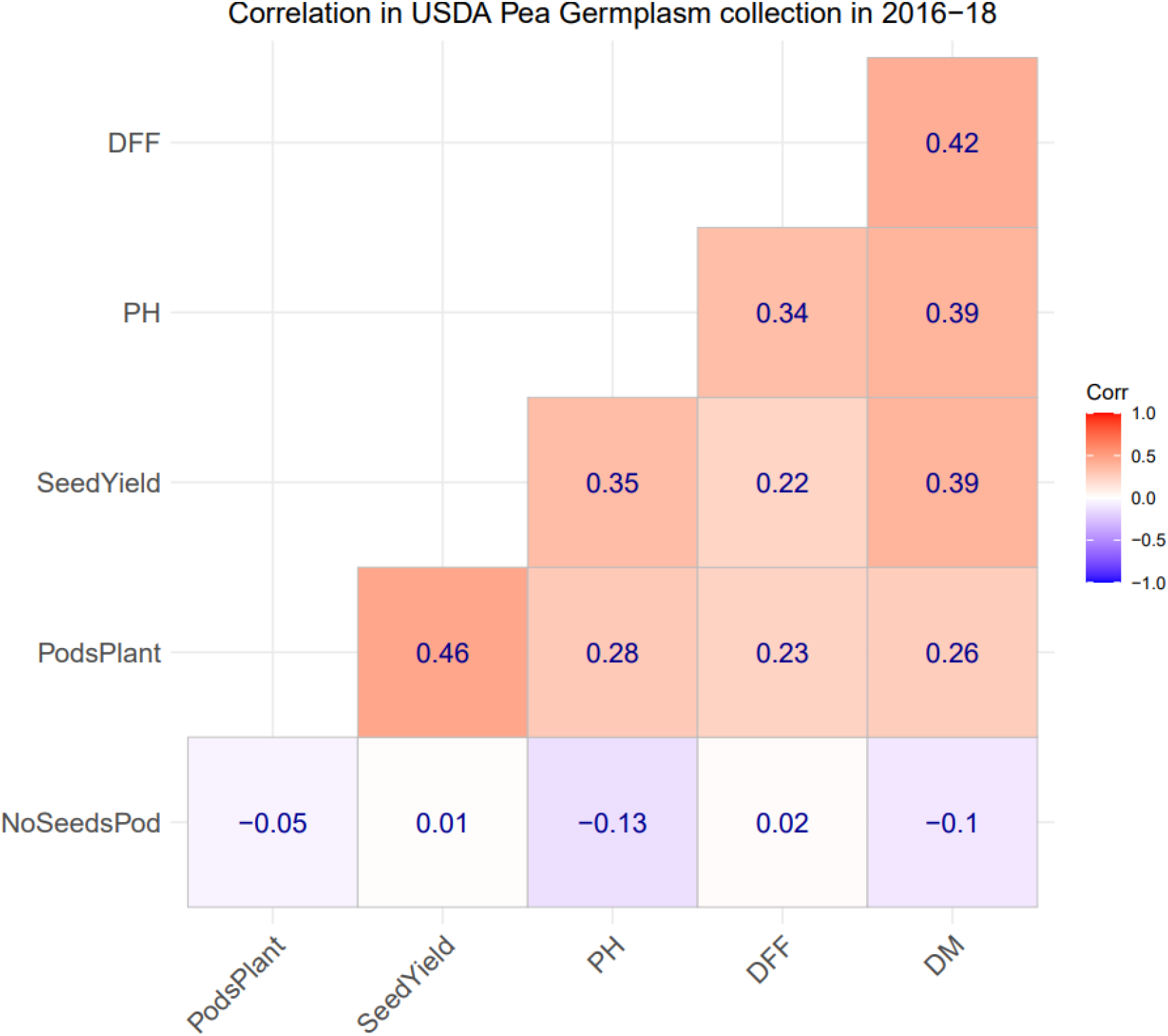
Phenotypic correlation among seed yield and agronomic traits evaluated in this study, DFF is days to first flowering, PH is plant height in cm, SeedYield is seed yield in kg ha^−1^, DM is the days to physiological maturity

**Supplementary Figure S2.**
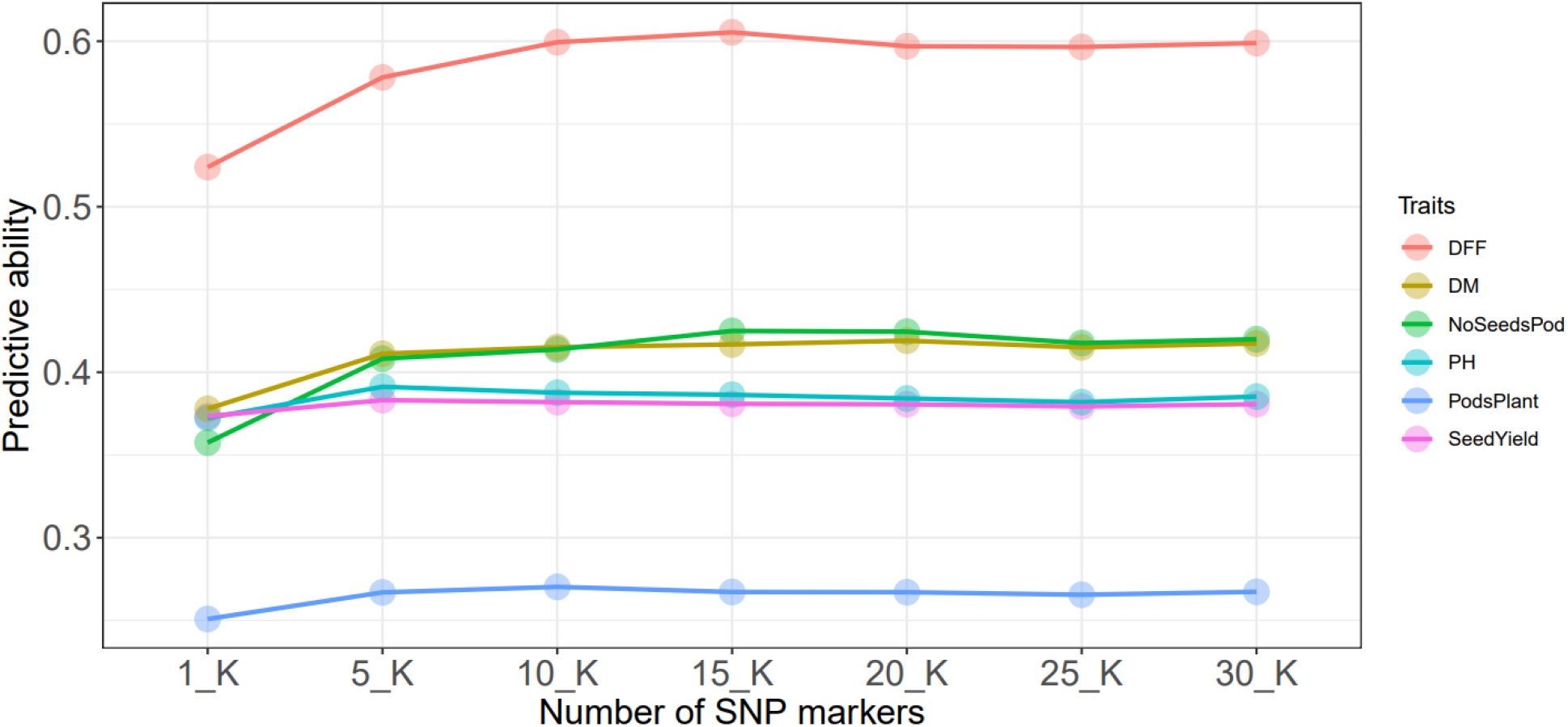
Predictive ability with increasing SNP markers RR-BLUP model, DFF is days to first flowering, DM, is days to physiological maturity, NoSeedsPod is number of seeds per pod, PH is plant height in cm, PodsPlant is pods per plant, SeedYield is seed yield in kg ha^−1^

**Supplementary Figure S3.**
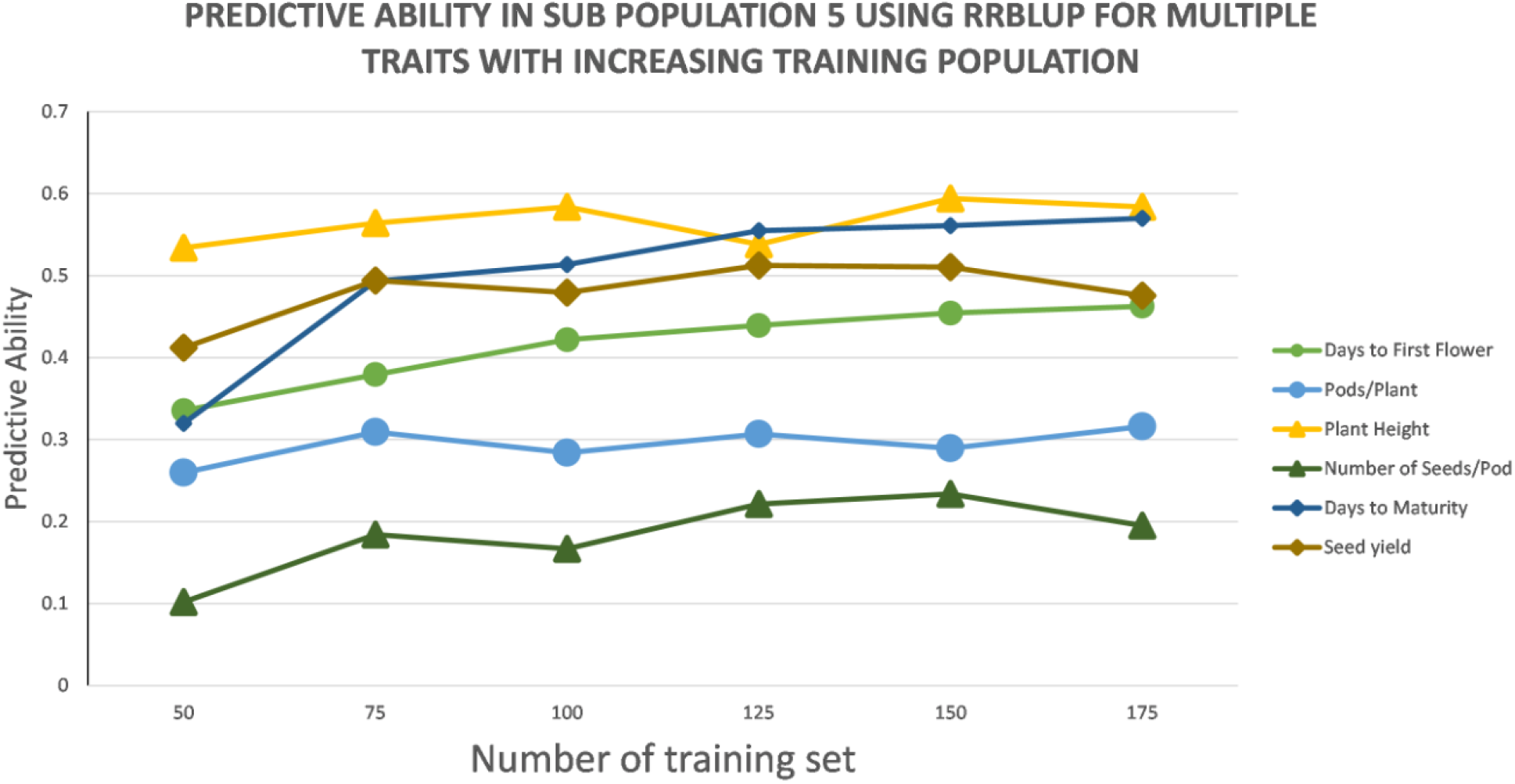
Predictive ability of subpopulation 5 with increasing training population

**Supplementary Figure S4.**
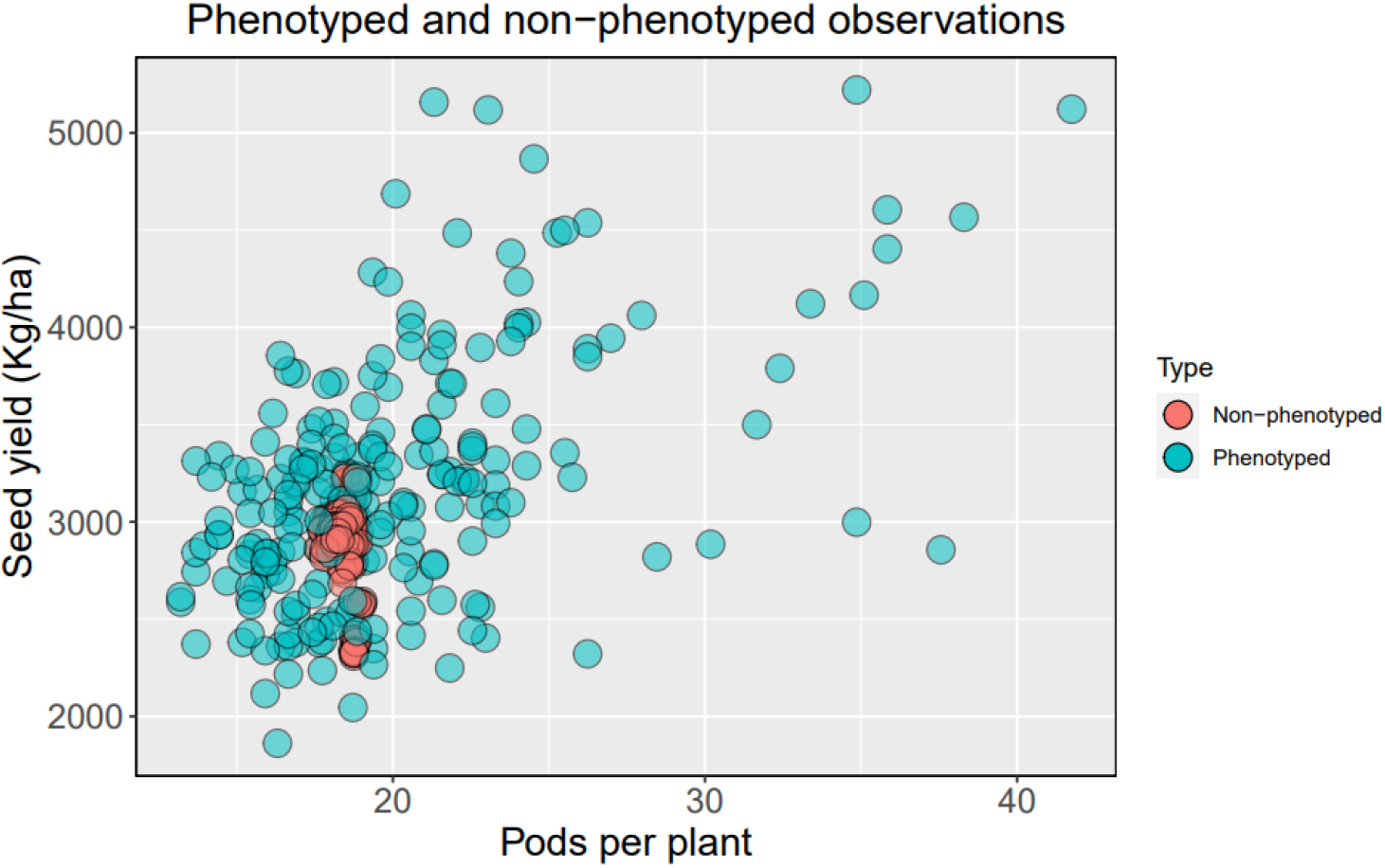
Distribution of phenotyped and predicted non-phenotyped accessions for seed yield and number of pods per plant in the USDA germplasm collections

**Supplementary Figure S5.**
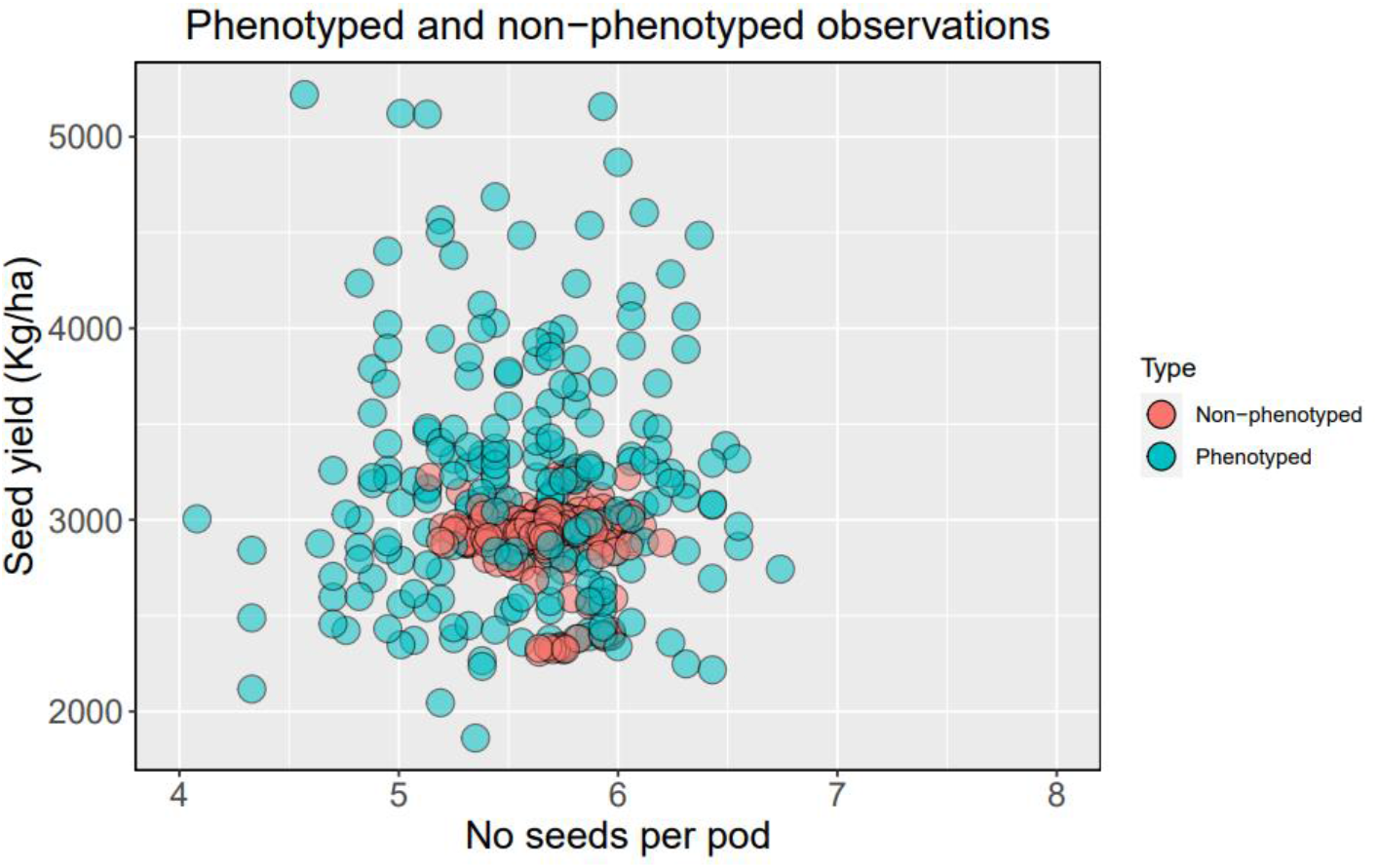
Distribution of phenotyped and predicted non-phenotyped accessions for seed yield and number of seeds per pod in the USDA germplasm collections

## References

Alexander, D.H., Novembre, J. and Lange, K., 2009. Fast model-based estimation of ancestry in unrelated individuals. Genome Research, 19(9), pp.1655–1664.

Annicchiarico, P., Nelson N., Meriem L., Imane Thami-Alami, Massimo R., and Luciano P. 2020. “Development and Proof-of-Concept Application of Genome-Enabled Selection for Pea Grain Yield under Severe Terminal Drought.” International Journal of Molecular Sciences 21 (7): 1–20. https://doi.org/10.3390/ijms21072414.

Annicchiarico, P., Nelson N., Luciano P., Massimo R., and Luigi R. 2019. “Pea Genomic Selection for Italian Environments.” BMC Genomics 20 (1): 1–18. https://doi.org/10.1186/s12864-019-5920-x.

Azodi, Christina B., Emily B., Andrew M., Mark R., Gustavo de los Campos, and Shin H. S. 2019. “Benchmarking Parametric and Machine Learning Models for Genomic Prediction of Complex Traits.” G3: Genes, Genomes, Genetics 9 (11): 3691–3702. https://doi.org/10.1534/g3.119.400498.

Bates, D., Martin M., Benjamin M. B., and Steven C. W. 2015. “Fitting Linear Mixed-Effects Models Using Lme4.” Journal of Statistical Software 67 (1). https://doi.org/10.18637/jss.v067.i01.

Bethke, Paul C., Dennis A. H., and Shelley H. J. 2019. “Potato Germplasm Enhancement Enters the Genomics Era,” 1–20.

Bradbury, P. J., Zhiwu Z., Dallas E. K., Terry M. C., Yogesh R., and Edward S. B. 2007. “TASSEL: Software for Association Mapping of Complex Traits in Diverse Samples.” Bioinformatics 23 (19): 2633–35. https://doi.org/10.1093/bioinformatics/btm308.

Breiman, L., 2001 Random Forests. Mach. Learn. 45: 5–32. https://doi.org/10.1023/A:1010933404324

Burstin, Judith, Pauline Salloignon, Marianne Chabert-Martinello, Jean Bernard Magnin-Robert, Mathieu Siol, Françoise Jacquin, Aurélie Chauveau, et al. 2015. “Genetic Diversity and Trait Genomic Prediction in a Pea Diversity Panel.” BMC Genomics 16 (1): 1–17. https://doi.org/10.1186/s12864-015-1266-1.

Cheng, Peng, William Holdsworth, Yu Ma, Clarice J. Coyne, Michael Mazourek, Michael A. Grusak, Sam Fuchs, and Rebecca J. McGee. 2015. “Association Mapping of Agronomic and Quality Traits in USDA Pea Single-Plant Collection.” Molecular Breeding 35 (2). https://doi.org/10.1007/s11032-015-0277-6.

Clark, Samuel A., John M. Hickey, Hans D. Daetwyler, and Julius H.J. van der Werf. 2012. “The Importance of Information on Relatives for the Prediction of Genomic Breeding Values and the Implications for the Makeup of Reference Data Sets in Livestock Breeding Schemes.” Genetics, Selection, Evolution : GSE 44 (1): 4. https://doi.org/10.1186/1297-9686-44-4.

Colombani, C., P. Croiseau, S. Fritz, F. Guillaume, A. Legarra, V. Ducrocq, and C. Robert-Granié. 2012. “A Comparison of Partial Least Squares (PLS) and Sparse PLS Regressions in Genomic Selection in French Dairy Cattle.” Journal of Dairy Science 95 (4): 2120–31. https://doi.org/10.3168/jds.2011-4647.

Covarrubias-Pazaran, Giovanny. 2016. “Genome-Assisted Prediction of Quantitative Traits Using the r Package Sommer.” PLoS ONE 11 (6): 1–15. https://doi.org/10.1371/journal.pone.0156744.

Coyne, C J, A F Brown, G M Timmerman-Vaughan, K E McPhee, and M A Grusak. 2005. “USDA-ARS Refined Pea Core Collection for 26 Quantitative Traits.” Pisum Genetics 37 (11): 1–4.

Cullis, Brian R., A. B. Smith, and N. E. Coombes. 2006. “On the Design of Early Generation Variety Trials with Correlated Data.” Journal of Agricultural, Biological, and Environmental Statistics 11 (4): 381–93. https://doi.org/10.1198/108571106X154443.

Crossa, José, Diego Jarquín, Jorge Franco, Paulino Pérez-Rodríguez, Juan Burgueño, Carolina Saint-Pierre, Prashant Vikram, et al. 2016. “Genomic Prediction of Gene Bank Wheat Landraces.” G3: Genes, Genomes, Genetics 6 (7): 1819–34. https://doi.org/10.1534/g3.116.029637.

Crossa, José, Paulino Pérez-rodríguez, Jaime Cuevas, Osval Montesinos-lópez, Diego Jarquín, Gustavo De Los Campos, Juan Burgueño, et al. 2017. “Genomic Selection in Plant Breeding : Methods, Models, and Perspectives.” Trends in Plant Science xx: 1–15. https://doi.org/10.1016/j.tplants.2017.08.011.

Danecek, P., Auton, A., Abecasis, G., Albers, C. A., Banks, E., DePristo, M. A., … Durbin, R. 2011. The variant call format and VCFtools. Bioinformatics, 27(15), 2156–2158. https://doi.org/10.1093/bioinformatics/btr330.

de los Campos, G., Hickey, J., Pong-Wong, R., Daetwyler, H. and M. Calus, 2013. Whole-genome regression and prediction methods applied to plant and animal breeding. Genetics 193: 327–345. https://doi.org/10.1534/genetics.112.143313.

de los Campos, Gustavo De, Daniel Gianola, Guilherme J.M. Rosa, Kent A. Weigel, and Jos Crossa. 2010. “Semi-Parametric Genomic-Enabled Prediction of Genetic Values Using Reproducing Kernel Hilbert Spaces Methods.” Genetics Research 92 (4): 295–308. https://doi.org/10.1017/S0016672310000285.

Elshire, R. J., Glaubitz, J. C., Sun, Q., Poland, J. A., Kawamoto, K., Buckler, E. S., & Mitchell, S. E. (2011). A robust, simple genotyping-by-sequencing (GBS) approach for high diversity species. PLoS ONE, 6(5), 1–10. https://doi.org/10.1371/journal.pone.0019379

Endelman, Jeffrey B. 2011. “Ridge Regression and Other Kernels for Genomic Selection with R Package RR-BLUP.” The Plant Genome 4 (3): 250–55. https://doi.org/10.3835/plantgenome2011.08.0024.

Facciolongo, Anna Maria, Giuseppe Rubino, Antonia Zarrilli, Arcangelo Vicenti, Marco Ragni, and Francesco Toteda. 2014. “Alternative Protein Sources in Lamb Feeding 1. Effects on Productive Performances, Carcass Characteristics and Energy and Protein Metabolism.” Progress in Nutrition 16 (2): 105–15.

Garrison, E., & Marth, G. 2012. Haplotype-based variant detection from short-read sequencing. ArXiv: 1207.3907 [q-Bio]. Retrieved from http://arxiv.org/abs/1207.3907.

Gaynor, R.C. 2015. GSwGBS: An R package genomic selection with genotyping-by-sequencing. Genomic selection for Kansas wheat. K-State Research Exchange, Manhattan, KS.

Gorjanc, Gregor, Janez Jenko, Sarah J Hearne, and John M Hickey. 2016. “Initiating Maize Pre-Breeding Programs Using Genomic Selection to Harness Polygenic Variation from Landrace Populations.” BMC Genomics 17 (1): 1–15. https://doi.org/10.1186/s12864-015-2345-z.

Guo, Jia, Sumit Pradhan, Dipendra Shahi, Jahangir Khan, Jordan Mcbreen, Guihua Bai, J. Paul Murphy, and Md Ali Babar. 2020. “Increased Prediction Accuracy Using Combined Genomic Information and Physiological Traits in A Soft Wheat Panel Evaluated in Multi-Environments.” Scientific Reports 10 (1): 1–12. https://doi.org/10.1038/s41598-020-63919-3.

Haile, Teketel A., Taryn Heidecker, Derek Wright, Sandesh Neupane, Larissa Ramsay, Albert Vandenberg, and Kirstin E. Bett. 2020. “Genomic Selection for Lentil Breeding: Empirical Evidence.” Plant Genome 13 (1): 1–15. https://doi.org/10.1002/tpg2.20002.

Habier, D, R L Fernando, and J C M Dekkers. 2007. “The Impact of Genetic Relationship Information on Genome-Assisted Breeding Values.” https://doi.org/10.1534/genetics.107.081190.

Hallauer, A R, M J Carena, and J B Miranda Fo. 2010. Hand Book of Plant Breeding: Quantitative genetics in maize breeding. 3rd ed. Springer, New York.

Hayes, B J, P J Bowman, A J Chamberlain, and M E Goddard. 2009. “Invited Review : Genomic Selection in Dairy Cattle : Progress and Challenges.” Journal of Dairy Science 92 (2): 433–43. https://doi.org/10.3168/jds.2008-1646.

Hickey, John M., Susanne Dreisigacker, Jose Crossa, Sarah Hearne, Raman Babu, Boddupalli M. Prasanna, Martin Grondona, et al. 2014. “Evaluation of Genomic Selection Training Population Designs and Genotyping Strategies in Plant Breeding Programs Using Simulation.” Crop Science 54 (4): 1476–88. https://doi.org/10.2135/cropsci2013.03.0195.

Holdsworth, William L., Elodie Gazave, Peng Cheng, James R. Myers, Michael A. Gore, Clarice J. Coyne, Rebecca J. McGee, and Michael Mazourek. 2017. “A Community Resource for Exploring and Utilizing Genetic Diversity in the USDA Pea Single Plant plus Collection.” Horticulture Research 4 (January). https://doi.org/10.1038/hortres.2017.17.

James, G., Witten, D., Hastie, T., Tibshirani, R. 2013. An Introduction to Statistical Learning: with Applications in R. Springer, New York. ISBN 978-1-4614-7138-7(eBook).

Jarquin, Diego, James Specht, and Aaron Lorenz. 2016. “Prospects of Genomic Prediction in the USDA Soybean Germplasm Collection: Historical Data Creates Robust Models for Enhancing Selection of Accessions.” G3: Genes, Genomes, Genetics 6 (8): 2329–41. https://doi.org/10.1534/g3.116.031443.

Kremling KAG, Diepenbrock CH, Gore MA, Buckler ES, Bandillo NB. 2019. Transcriptome-Wide Association Supplements Genome-Wide Association in Zea mays. G3. 9:3023–3033.

Kreplak, J., Madoui, M.A., Cápal, P., Novák, P., Labadie, K., et al, 2019. A reference genome for pea provides insight into legume genome evolution. Nature Genetics, 51(9), pp.1411–1422.

Li H. and Durbin R. 2009. Fast and accurate short read alignment with Burrows-Wheeler Transform. Bioinformatics, 25:1754–60).

Li, H., Handsaker, B., Wysoker, A., Fennell, T., Ruan, J., Homer, N., Marth, G., Abecasis, G. and Durbin, R., 2009. The sequence alignment/map format and SAMtools. Bioinformatics, 25(16), pp.2078–2079.

Li, Huihui, Awais Rasheed, Lee T. Hickey, and Zhonghu He. 2018. “Fast-Forwarding Genetic Gain.” Trends in Plant Science 23 (3): 184–86. https://doi.org/10.1016/j.tplants.2018.01.007.

Li, Xin, Tingting Guo, Qi Mu, Xianran Li, and Jianming Yu. 2018. “Genomic and Environmental Determinants and Their Interplay Underlying Phenotypic Plasticity.” Proceedings of the National Academy of Sciences of the United States of America 115 (26): 6679–84. https://doi.org/10.1073/pnas.1718326115.

Liu, Z., F. Seefried, F. Reinhardt, S. Rensing, G. Thaller et al., 2011 Impacts of both reference population size and inclusion of a residual polygenic effect on the accuracy of genomic prediction. Genet. Sel. Evol. 43: 19. https://doi.org/10.1186/1297-9686-43-19.

Longin, C. Friedrich H., and Jochen C. Reif. 2014. “Redesigning the Exploitation of Wheat Genetic Resources.” Trends in Plant Science 19 (10): 631–36. https://doi.org/10.1016/j.tplants.2014.06.012.

Lorenz, A. J. & Smith, K. P. 2015. Adding genetically distant individuals to training populations reduces genomic prediction accuracy in barley. Crop Sci. 55, 2657–2667.

Mascher, Martin, Mona Schreiber, Uwe Scholz, Andreas Graner, Jochen C. Reif, and Nils Stein. 2019. “Genebank Genomics Bridges the Gap between the Conservation of Crop Diversity and Plant Breeding.” Nature Genetics 51 (7): 1076–81. https://doi.org/10.1038/s41588-019-0443-6.

Meuwissen, T H E, B J Hayes, and M E Goddard. 2001. “Prediction of Total Genetic Value Using Genome-Wide Dense Marker Maps.”

Mevik, B.-H., & Wehrens, R. (2007). The pls Package: Principal Component and Partial Least Squares Regression in R. Journal of Statistical Software, 18(2), 1–23. https://doi.org/10.18637/jss.v018.i02.

Money, Daniel, Kyle Gardner, Zoë Migicovsky, Heidi Schwaninger, Gan Yuan Zhong, and Sean Myles. 2015. “LinkImpute: Fast and Accurate Genotype Imputation for Nonmodel Organisms.” G3: Genes, Genomes, Genetics 5 (11): 2383–90. https://doi.org/10.1534/g3.115.021667.

Mudryj, Adriana N., Nancy Yu, and Harold M. Aukema. 2014. “Nutritional and Health Benefits of Pulses.” Applied Physiology, Nutrition and Metabolism 39 (11): 1197–1204. https://doi.org/10.1139/apnm-2013-0557.

Norman, Adam, Julian Taylor, James Edwards, and Haydn Kuchel. 2018. “Optimising Genomic Selection in Wheat: Effect of Marker Density, Population Size and Population Structure on Prediction Accuracy.” G3: Genes, Genomes, Genetics 8 (9): 2889–99. https://doi.org/10.1534/g3.118.200311.

Pérez, Paulino, and Gustavo De Los Campos. 2014. “Genome-Wide Regression and Prediction with the BGLR Statistical Package.” Genetics 198 (2): 483–95. https://doi.org/10.1534/genetics.114.164442.

R Core Team. 2020. R: A language and environment for statistical computing. R Foundation for Statistical Computing, Vienna, Austria. https://www.R-project.org/.

Riedelsheimer, Christian, Yariv Brotman, Michaël Méret, Albrecht E. Melchinger, and Lothar Willmitzer. 2013. “The Maize Leaf Lipidome Shows Multilevel Genetic Control and High Predictive Value for Agronomic Traits.” Scientific Reports 3: 1–7. https://doi.org/10.1038/srep02479.

Rutkoski, J., R. P. Singh, J. Huerta‐Espino, S. Bhavani, J. Poland, J. L. Jannink, and M. E. Sorrells. 2015. “Efficient Use of Historical Data for Genomic Selection: A Case Study of Stem Rust Resistance in Wheat.” The Plant Genome 8 (1): 1–10. https://doi.org/10.3835/plantgenome2014.09.0046.

Simson, C. J. & Hannan, R. M. 1995. “Development and Use of Core Subsets of Cool-Season Food Legume Germplasm Collections.” HortScience 30: 907.

Spindel, Jennifer, Hasina Begum, Deniz Akdemir, Parminder Virk, Bertrand Collard, Edilberto Redoña, Gary Atlin, Jean Luc Jannink, and Susan R. McCouch. 2015. “Genomic Selection and Association Mapping in Rice (*Oryza sativa*): Effect of Trait Genetic Architecture, Training Population Composition, Marker Number and Statistical Model on Accuracy of Rice Genomic Selection in Elite, Tropical Rice Breeding Lines.” PLoS Genetics 11 (2): 1–25. https://doi.org/10.1371/journal.pgen.1004982.

Tayeh, Nadim, Anthony Klein, Marie Christine Le Paslier, Françoise Jacquin, Hervé Houtin, Céline Rond, Marianne Chabert-Martinello, et al. 2015. “Genomic Prediction in Pea: Effect of Marker Density and Training Population Size and Composition on Prediction Accuracy.” Frontiers in Plant Science 6 (NOVEMBER): 1–11. https://doi.org/10.3389/fpls.2015.00941.

USDA. 2020. “United States Acreage,” 1–50. https://www.nass.usda.gov/Publications/Todays_Reports/reports/acrg0620.pdf.

Valluru, R., Gazave, E. E., Fernandes, S. B., Ferguson, J. N., Lozano, R., Hirannaiah, P., … Bandillo, N. 2019. Deleterious mutation burden and its association with complex traits in sorghum (*Sorghum bicolor*). Genetics, 211(3), 1075 LP – 1087.

Vandemark, G J, M Brick, J M Osorno, D J Kelly & C A Urrea. 2014. Edible grain legumes. In S Smith, B Diers, J. Speecht, & B Carver (Eds.), Yield Grains in major U.S. field crops (pp.87–123). Madison, WI: CSSA. https://doi.org/10.3390/cli6020041.

VanRaden, P. M. 2008. “Efficient Methods to Compute Genomic Predictions.” Journal of Dairy Science 91 (11): 4414–23. https://doi.org/10.3168/jds.2007-0980.

Wickham H (2016). ggplot2: Elegant Graphics for Data Analysis. Springer-Verlag New York. ISBN 978-3-319-24277-4.

Yu, Xiaoqing, Samuel Leiboff, Xianran Li, Tingting Guo, Natalie Ronning, Xiaoyu Zhang, Gary J. Muehlbauer, et al. 2020. “Genomic Prediction of Maize Microphenotypes Provides Insights for Optimizing Selection and Mining Diversity.” Plant Biotechnology Journal, 2456–65. https://doi.org/10.1111/pbi.13420.

Yu, Xiaoqing, Xianran Li, Tingting Guo, Chengsong Zhu, Yuye Wu, Sharon E. Mitchell, Kraig L. Roozeboom, et al. 2016. “Genomic Prediction Contributing to a Promising Global Strategy to Turbocharge Gene Banks.” Nature Plants 2 (October). https://doi.org/10.1038/nplants.2016.150.

